# Frataxin deficiency in the astrocytes drives neurocognitive impairment in sickle cell disease mice

**DOI:** 10.1101/2024.12.20.629760

**Authors:** Enrico M. Novelli, Shane C Lenhart, Lesley M. Foley, Nandinii Sekar, Paritosh Mondal, Hong Wang, T. Kevin Hitchens, Samit Ghosh, Stephen Y. Chan, Xiaoming Hu, Rimi Hazra

## Abstract

Individuals with sickle cell disease (SCD) suffer from a high burden of cerebrovascular lesions and cognitive impairment that vastly impact quality of life. Cerebrovascular lesions are characterized by microstructural neuroaxonal damage, but their pathogenesis has not been fully elucidated. Herein, we report that SCD mice (SS) have reduced expression of frataxin (FXN), a mitochondrial protein, in their astrocytes compared to control (AA) mice. Next, we generated chimeric mice with SS bone marrow and astrocyte-specific deletion of FXN (SS^FXN-KO^). Ex-vivo diffusion tensor magnetic resonance imaging and immunohistopathology of the brain showed that the SS^FXN-KO^ mice have increased white matter neuroaxonal damage compared to the SS bone marrow chimera mice with wild-type FXN expression (SS^FXN-WT^). The SS^FXN-KO^ mice also displayed poorer cognitive function as measured by the functional Y-maze and novel object recognition tests. Pharmacological induction of FXN by administration of insulin growth factor-1 improved cognitive function in the SS^FXN-KO^ mice. Overall, our data demonstrate that FXN is a critical factor regulating neuroaxonal health and cognitive function in SCD mice. FXN may therefore be a novel pharmacologic target to prevent cerebrovascular complications in SCD.

## Main Text

Individuals with sickle cell disease (SCD) suffer from a high burden of cerebrovascular lesions and cognitive impairment that vastly impact quality of life. About 40% of children with SCD develop infarcts, while about 55% of them show delays in early markers of cognition and expressive language. Difficulties in processing speed, working memory, adaptation and behavior continue to progress across the lifespan. Cerebrovascular lesions are primarily diagnosed by T2-weighted magnetic resonance imaging (MRI) in combination with diffusion tensor imaging (DTI), which have revealed widespread white matter abnormalities with lower fractional anisotropy (FA) in SCD. A high prevalence of cognitive impairment is associated with white matter injury in SCD (1). Astrocytes interconnect cerebral microvasculature and neurons and maintain neuronal integrity by regulating cerebral blood flow, calcium oscillation, and preserving plasticity and memory function (2). Microstructural white matter injury is associated with activation of astrocytes in transgenic SCD (SS) mice that simultaneously displayed poorer cognitive function compared to non-SCD control mice (3). Herein, we investigated a potential mechanism by which activated astrocytes mediate cerebrovascular pathology in the SS mice.

Frataxin (FXN) regulates iron-sulfur biogenesis, and its deficiency is associated with deregulation of calcium signaling, and disruption of mitochondrial function in the cerebrovascular endothelium and astrocytes, regulating neurocognitive dysfunction (4). We tested whether FXN deficiency in the astrocytes is responsible for neurocognitive impairment in SS mice.

### Astrocytic FXN is critical to maintain neuroaxonal integrity in SCD mice

In SS mice, the widespread abnormalities in the brain identified by MRI and DTI are directly linked to histopathological evidence of demyelinated neuroaxonal damage, and are associated with astrocyte activation identified by higher expression of glial fibrillar acidic protein (GFAP) (3). We discovered that in SS mice brains, expression of FXN was significantly downregulated in GFAP+ astrocytes compared to their AA littermates **(Figure 1A-B)**. We postulate that inhibition of FXN activates astrocytes, which in turn exacerbates neurocognitive impairment in SCD. To test this hypothesis, we generated tamoxifen inducible astrocyte-specific FXN knockout mice (FXN-KO) by crossing FXN-floxed mice with *Aldh1l1*-Cre mice. The FXN-KO mice had drastically reduced astrocytic FXN expression following tamoxifen injection **(Supplementary Figure 1A-C)**. We created SCD bone marrow chimera mice with the SCD phenotype on the FXN-KO (SS^FXN-KO^) and FXN-WT (SS^FXN-WT^) background **(Supplementary Figure 2A-B)**. After successful engraftment (8-12 weeks following BMT), the SS^FXN-KO^ and SS^FXN-WT^ mice were anemic like the donor SS mice with low hematocrit and high reticulocyte counts **(Supplementary Table 1)**. The SS^FXN-KO^ mice were injected with tamoxifen to knock down astrocytic FXN **(Supplementary Figure 2C)**. The brains of the chimeric mice were scanned by MRI, followed by histopathology to detect microstructural changes in the white matter. From the ex-vivo DTI of the brains, we generated diffusion-encoded color (DEC) maps of the SS^FXN-KO^ and SS^FXN-WT^ mice (**Figure 1C)**. The corpus callosum (CC) and the external capsule (EC) areas of the brain showed significantly reduced FA, a white matter injury indicator based on water diffusion directionality, in the SS^FXN-KO^ mice compared to the SS^FXN-WT^ mice **(Figure 1D)**. The multidirectional diffusivity, such as axial diffusivity (AD) and radial diffusivity (RD) are considered as surrogate DTI markers for axonal and myelin damage respectively. The SS^FXN-KO^ mice displayed a substantial reduction in AD **(Figure 1E)**, while the reduction in RD was less prominent **(Figure 1F)**. An increased ratio of SMI32/MBP is a marker of neuroaxonal damage and we found that the brain tissue showed substantial elevation in non-phosphorylated neurofilament H (SMI32) with a concomitant decrease in myelin basic protein (MBP) in SS^FXN-KO^ mice **(Figure 1G-H)**. Moreover, we found an increased number of GFAP+ activated astrocytes in SS^FXN-KO^ mice brain **(Figure 1I-J)**. These data suggest that FXN is a critical factor that regulates neuroaxonal health and that its deficiency phenocopies widespread reduction in FA and increased RD, characteristics of microstructural neuroaxonal damage in SCD patients.

**Figure 1.**
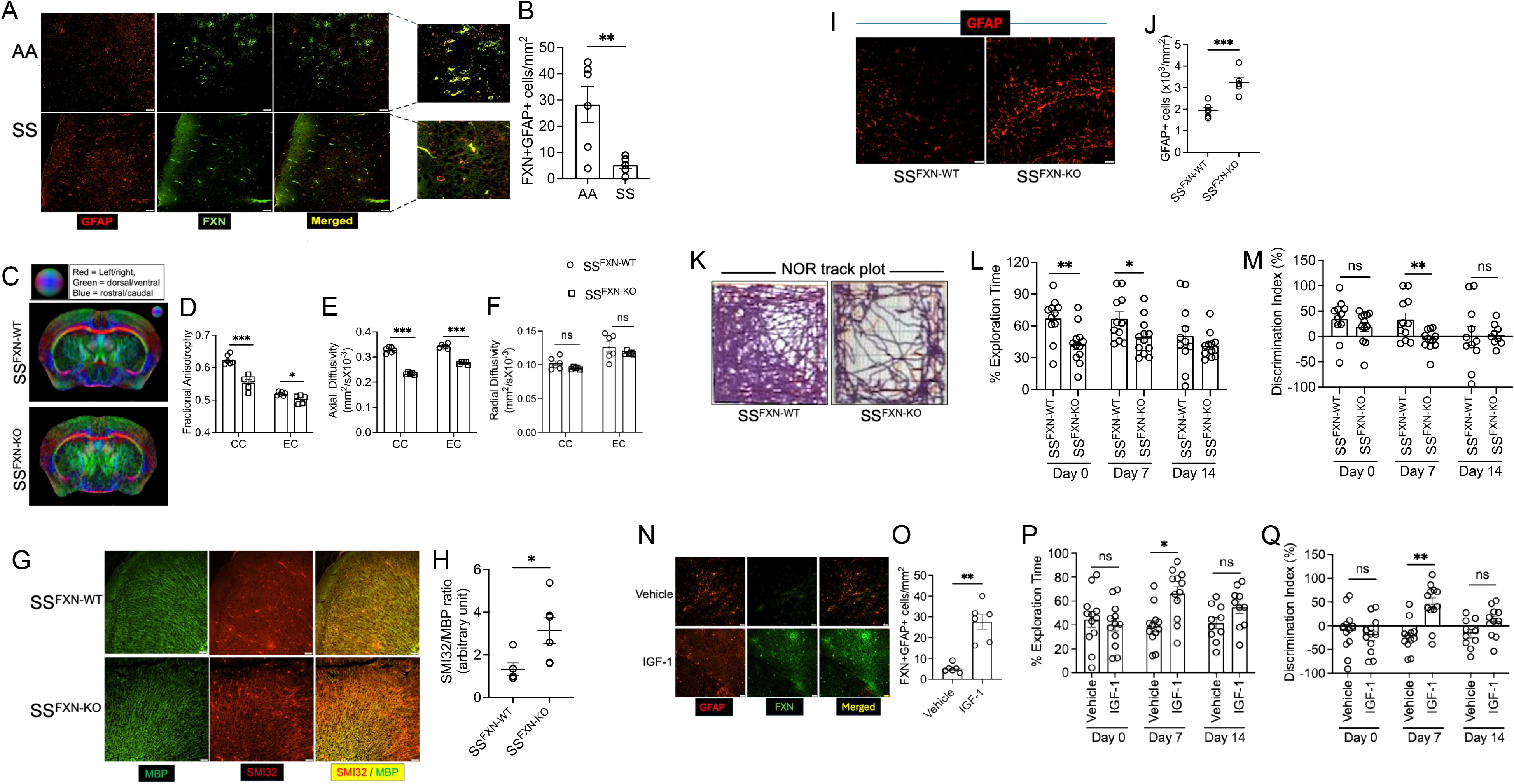
Frataxin deficiency is linked to the reduced neuroaxonal integrity and cognitive impairment in sickle mice. **A)** Representative immunofluorescence imaging showing decreased frataxin expression in GFAP+ astrocytes of SS mice compared to AA mice (scale bar = 50μm). **(B)** Quantitation of FXN+GFAP+ cells in AA and SS mouse brain tissue (n=6). **(C)** Representative DEC maps for SS^FXN-WT^ and SS^FXN-KO^ mice. Comparison of scalar diffusion parameters, **(D)** Fractional anisotropy, **(E)** axial diffusivity, and **(F)** radial diffusivity, for the external capsule (EC) and corpus callosum (CC) ROIs between SS^FXN-WT^ and SS^FXN-KO^ mice (n=6). **(G-H)** Immunofluorescence staining showing elevated SMI32/MBP staining intensity in SS^FXN-KO^ mice compared to SS^FXN-WT^ mice (n=6; scale bar = 50μm). **(I-J)** Increased astrocyte activation marked by amplified GFAP+ staining in SS^FXN-KO^ mice (n=6; scale bar = 50μm). **(K)** Representative track plot from NOR testing showed halted movement of SS^FXN-KO^ mice. **(L-M)** Both SS^FXNWT^ and SS^FXN-KO^ mice were tested for NOR for 3 days on weekly interval. Reduced exploration time (L) and discrimination index (M) were evident in SS^FXN-KO^ mice (n=11-12). **(N)** The SS mice treated with recombinant IGF-1 displayed increased FXN staining in their brain tissue compared to vehicle treated mice (scale bar = 50μm). **(O)** Quantitation of FXN+GFAP+ cells in IGF-1 and vehicle treated mice (n=6). **(P-Q)** The SS mice treated with IGF-1 showed improved cognitive function in NOR testing compared to vehicle treated mice (n=11-12). ns, non-significant; *p<0.05, **p<0.01, ***p<0.001 (unpaired t-test).

### Absence of FXN worsens while its induction improves neurocognitive responses in SCD mice

Since neuroaxonal damage in SCD is associated with impaired cognitive responses in both SCD mice and humans (3, 5), we determined whether deletion of astrocytic FXN leads to poor cognitive function in SCD mice. We performed novel object recognition (NOR) test, a behavioral non-spatial memory test, and the Y-maze test that identifies the inclination of the mice to exploring new environments in our experimental mice. The trajectory of animal movement showed a break in the continuity in absence of astrocytic FXN **(Figure 1K)**. The SS^FXN-KO^ mice spent significantly less time exploring the novel object than the SS^FXN-WT^ mice during the first two measurements within a 7-day interval (day 0 and 7) **(Figure 1L)**. The discrimination index, differentiating the time spent exploring the familiar object and the time spent with the novel object, was significantly lower in the SS^FXN-KO^ mice **(Figure 1M)**. In Y-maze, the frequency of spontaneous alternations — a measure of spatial working memory — indicates the tendency of the mice to alternate the exploration of different arms of a Y-shaped maze. We found that the SS^FXN-KO^ mice had reduced spontaneous alternations between the three arms of the maze compared to the SS^FXN-WT^ mice, which reached significance on the day 7 measurement **(Supplementary Figure 3A)**. However, the amount of time spent in a new arm of the Y-maze was unaltered **(Supplementary Figure 3B)**.

Prophylaxis with human recombinant insulin growth factor-1 (IGF-1) can stimulate FXN in neurons and astrocytes (6). We found pretreatment with IGF-1 considerably increased FXN expression in the astrocytes of the SS mice **(Figure 1N-O)**. We tested the cognitive responses of the SS mice treated with vehicle or IGF-1. Improved exploration time and discrimination index in the IGF-1 treated SS mice indicated that enhanced expression of FXN ameliorated cognitive function **(Figure 1P-Q)**. However, IGF-1 did not alter the rate of spontaneous alternations, or the time spent in a new arm of the Y-maze **(Supplementary Figure 3C-D)**.

In SCD, sustained chronic hemolytic stress, anemia and cerebral hypoxia lead to a compensatory increase in cerebral blood flow and hyperemia at baseline. Astrocytic mitochondria play a key role in the regulation of cerebral blood flow by altering the calcium dynamics within the astrocytic microdomains of the end feet that surround the cerebral microvasculature. Moreover, inhibition of FXN by hypoxia is linked to increase in vasoconstrictor production in endothelial cells. Severe hypoxia may, however, downregulate FXN and alter mitochondrial bioenergetics in ensheathing astrocytic end feet resulting in an increase in cerebral vasoconstriction. This may in turn abrogate the brain’s compensatory response to anemia and lead to neurocognitive dysfunction. Overall, our studies introduce astrocytic FXN as a critical component that may regulate cognitive function in SCD by preserving neuroaxonal integrity. Induction of FXN may be explored as a therapeutic strategy to target cerebrovascular complications in SCD.

## Acknowledgements

This work was supported by National Institutes of Health (NIH), *National Institute of Neurological Disorders and Stroke grant* 1R21NS131634-01A1 (RH), *National Institute of Diabetes and Digestive and Kidney* Diseases (NIDDK) grants R01DK124426 (SG), R01DK132145 (SG), and a P3HVB grant from the Hemophilia Center of Western Pennsylvania and Vitalant (RH). This work was supported by NIH grants R01 HL124021, HL122596, HL151228 (to S.Y.C.). *This work used Advanced Imaging Center (RRID:SCR_025139), a core research facility partially supported by the University of Pittsburgh and the office of the Senior Vice Chancellor for Health Sciences*.

## Authorship contributions

EMN critically reviewed the data and the manuscript, SCL generated the targeted frataxin knockout mice and performed the BMT experiments, NS performed the immunofluorescence staining. LMF and TKH performed, analyzed and interpreted DTI data, PM performed flow cytometry experiments, HW conducted statistical analysis, SG and SYC reviewed the manuscript, XU interpreted cognitive data and reviewed the manuscript, RH conceived and designed the study, performed experiments, analyzed and interpreted data, and wrote the manuscript with consultation and contribution from all coauthors.

## Data sharing

For original data, please contact

