## Supplemental figures and methods for "Frataxin deficiency in the astrocytes drives neurocognitive impairment in sickle cell disease mice"

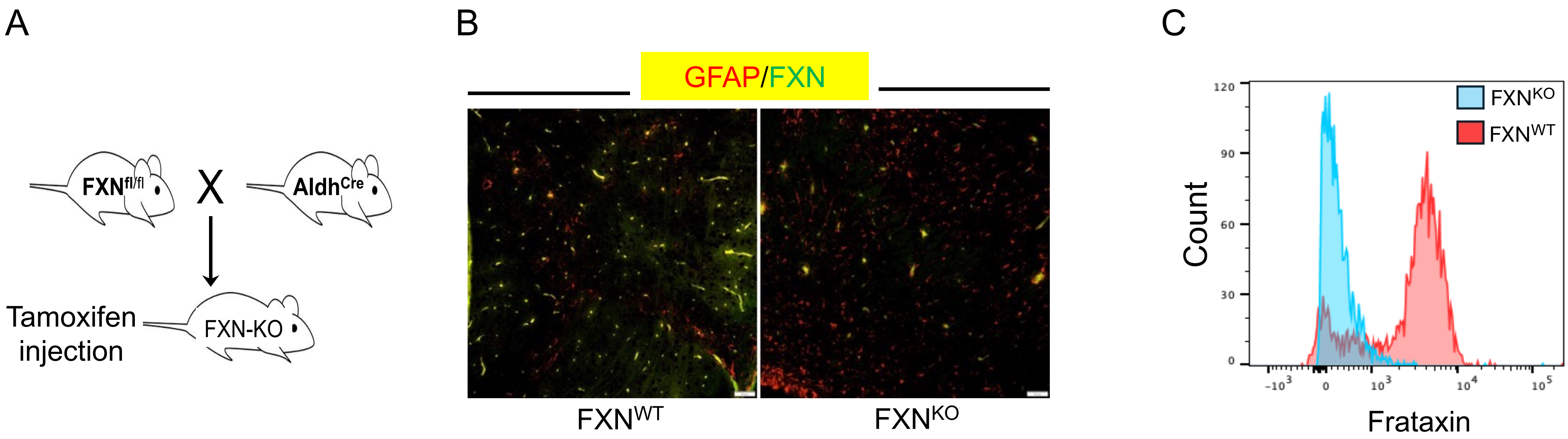


Supplementary Figure 1

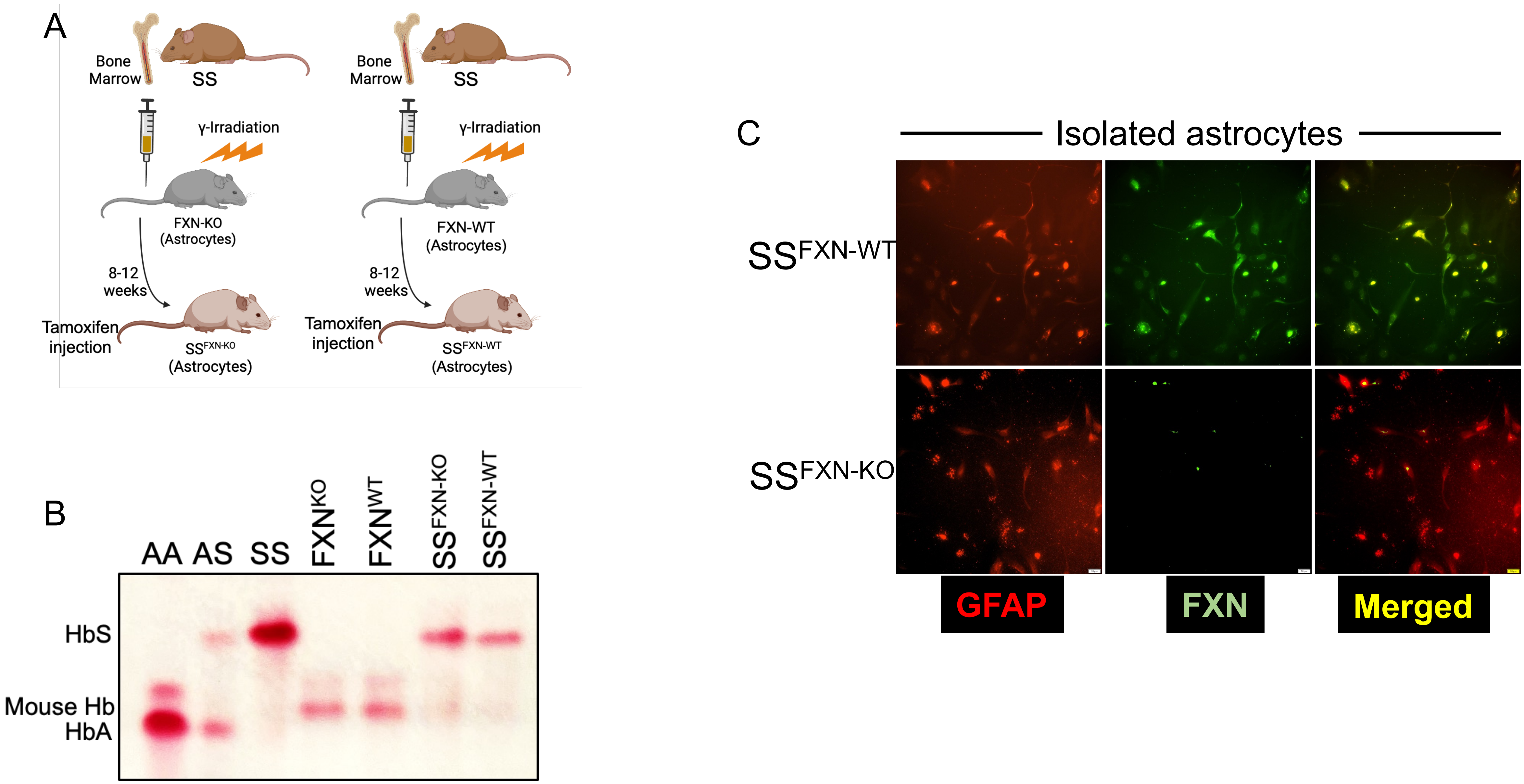


Supplementary Figure 2

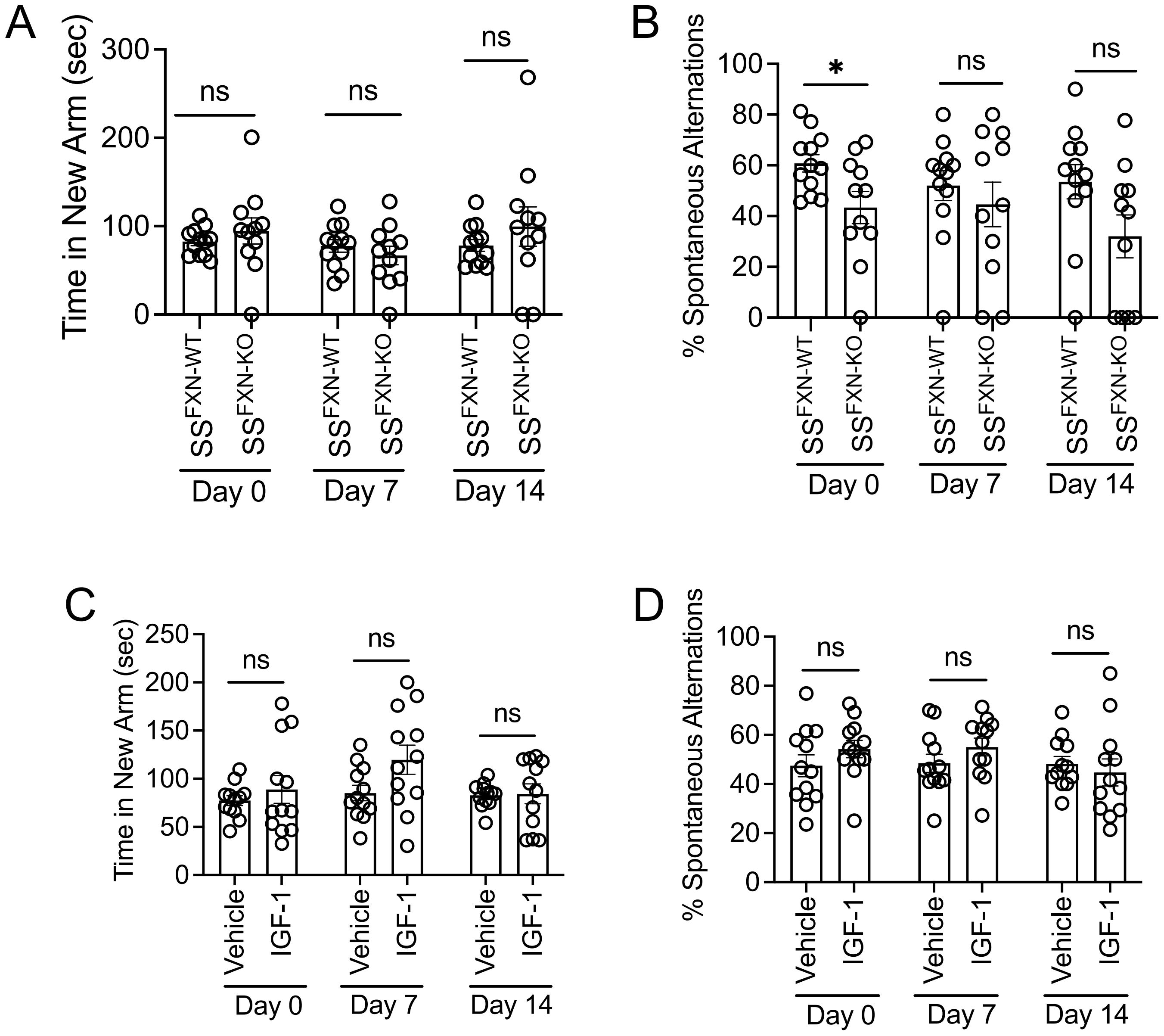


Supplementary Figure 3

| | SS | $FXN^{-WT}$ | $FXN^{-KO}$ | $SS^{FXN-KO}$ | $SS^{FXN-WT}$ |
|---|---|---|---|---|---|
| Hb (g/dl) | 6.9 ± 1.6 | 13.4 ± 0.33 | 12.1 ± 0.13 | 9.2 ± 0.51 | 8.3 ± 0.51 |
| HCT (%) | 30.3 ± 2.9 | 42.37 ± 1.08 | 43.21 ± 1.09 | 34.61 ± 2.39 | 33.51 ± 1.80 |
| Retics (%) | 59.6 ± 5.2 | 7.89 ± 1.06 | 10.29 ± 3.01 | 51.74 ± 7.19 | 43.88 ± 5.53 |
| WBC ($10^3$/µl) | 21.2 ± 8.1 | 6.18 ± 1.18 | 8.37 ± 0.94 | 19.6 ± 2.98 | 27.5 ± 3.84 |
| RBC($10^6$/µl) | 6.2 ± 2.13 | 8.71 ± 1.62 | 7.27 ± 1.38 | 7.02 ± 2.18 | 6.36 ± 1.08 |
| Platelet($10^3$/µl) | 398.2 ± 47.2 | 499.68 ± 98.1 | 378.23 ± 28.1 | 451.37 ± 71.2 | 415.57 ± 90.6 |

Hematological characteristics of donor, recipient and sickle bone marrow chimera mice as indicated. Data represented as mean ± SEM.

Supplementary Table 1

### Supplementary Figure Legends

**Supplementary Figure 1. Generation of mouse strain with targeted deletion of frataxin in astrocytes. (A)** General breeding schema between $FXN^{fl/fl}$ and $Aldh^{Cre}$ mice. **(B)** Representative cerebral tissue sections from $FXN^{WT}$ and $FXN^{KO}$ mice showing coexpression of FXN and GFAP (astrocyte marker) (scale bar=50 μm). **(C)** Flow analysis showing lack of FXN expression in the isolated astrocytes from the indicated group of mice.

**Supplementary Figure 2. Astrocyte specific FXN expression in $SS^{FXN\text{-}KO}$ and $SS^{FXN\text{-}WT}$. (A)** Schema showing generation of $SS^{FXN\text{-}KO}$ and $SS^{FXN\text{-}WT}$ mice. **(B)** Hemoglobin (Hb) gel electrophoresis showing expression of sickle Hb (HbS) in $SS^{FXN\text{-}KO}$ and $SS^{FXN\text{-}WT}$ mice indicating engraftment of sickle hemoglobin in $FXN^{KO}$ and $FXN^{WT}$ mice following sickle bone marrow transplantation. **(C)** Representative immunofluorescence images showing expression of FXN in GFAP+ astrocytes in $SS^{FXN\text{-}KO}$ and $SS^{FXN\text{-}WT}$ mice.

**Supplementary Figure 3. Changes in spatial working memory in sickle mice in absence or presence of frataxin. (A-B)** The $SS^{FXNWT}$ and $SS^{FXN\text{-}KO}$ mice were used for Y-maze behavioural testing to measure the time spent in a new arm of the maze (A), and spontaneous alternations (B) indicating percent frequency of the mice moving between two different arms in a Y-maze. **(C-D)** Cognitive activity of the IGF-1 treated SS mice in Y-maze testing (n=11-12).

### Supplementary Methods

#### *Insulin growth factor-1 Treatment*

Recombinant human IGF-1 (50 μg/kg, ab9573) or vehicle (phosphate buffer saline, PBS) were injected subcutaneously for 5 days.

#### *Mice and Bone marrow transplantation*

The University of Pittsburgh Institutional Animal Care and Use Committee (22010095) approved all mouse studies. Both male and female mice were used. The AA and SS mice were 12-14 weeks old at the time of the experiments. Knock-in Townes' SCD mice (#013071) [1] expressing human-Hb $\beta^S$ (SS) and control mice with human-Hb $\beta^A$ (AA) were purchased from Jackson Laboratories and bred in the University of Pittsburgh vivarium. Mouse genotypes were confirmed either by PCR or Hb gel electrophoresis. We bred floxed frataxin (FXN) mice ($FXN^{fl/fl}$; C57BL/6J-*Fxn*$^{em2Lutzy}$/J; #028520) with Aldh1l1-Cre/ERT2 BAC mice (B6N.FVB-Tg(Aldh1l1-cre/ERT2)1Khakh/J; #031008) to generate tamoxifen-inducible deletion of FXN specifically in *Aldh1l1*-expressing astrocytes ($FXN^{KO}$). The $FXN^{fl/fl}$ mice (referred herein as $FXN^{WT}$) (6-8 weeks of age, both sexes) were maintained on acidified drinking water for 7 days and subjected to one dose of 1200 rads irradiation. Irradiated mice were transplanted with $5x10^6$ whole bone marrow cells harvested from SS mouse donors. The transplanted mice (designated as $SS^{FXN-KO}$ and $SS^{FXN-WT}$) were maintained on medicated (Neomycin: 0.5mg/ml; Polymyxin B: 0.0125 mg/ml) water for one week. Mice were phlebotomized by retro orbital bleeding using a capillary tube internally coated with heparin/EDTA anticoagulant. Blood samples were collected from transplanted mice 8-weeks post-transplant for complete blood count (CBC) using

HemaTrue hematology analyzer (Heska). To assess reticulocyte count using flow cytometry. Percent reticulocytes were determined by flow cytometric analysis of thiazole orange stained whole blood samples [2] and analyzing data using a FACSDIVA software. Hb-gel electrophoresis was performed to identify the presence of HbS on the transplanted sickle bone marrow chimera mice.

***Diffusion tensor imaging (DTI)***

Ex-vivo DTI was performed using a Bruker AV3HD 11.7 Tesla/89 mm vertical-bore microimaging system equipped with a Micro2.5 gradient set, a 20 mm quadrature RF resonator and ParaVision 6.0.1 (Bruker Biospin, Billerica, MA). DTI data were collected using a multislice spin-echo sequence with 5 A0 images and 30 non-colinear diffusion-weighted images with the following parameters: TE/TR 22/2800 ms, 2 averages, 160 × 160 matrix, 16 × 16 mm field of view, 25 slices, 0.5 mm slice thickness, b-value = 3000 s/mm$^2$, and $\Delta/\delta$ = 11.0/5.0 ms. DTI data were analyzed with DSI Studio (http://dsi-studio.labsolver.org/) [3]. ROIs were drawn manually encompassing the corpus callosum (CC) or external capsule (EC) in the left and right hemispheres to determine mean fractional anisotropy (FA), axial diffusivity (AD) and radial diffusivity (RD) for $SS^{FXNWT}$ and $SS^{FXNKO}$ mice.

***Immunofluorescence***

Following DTI scanning, the whole brains were isolated. The brain tissue from individual mice were cut into 3 mm slices. The 2$^{nd}$ slice from the anterior half of the brain was processed for paraformaldehyde-fixed, paraffin embedded sectioning (4 $\mu$m). A total of 6 tissue sections per brain per mouse were taken from the 2$^{nd}$ slice for immunofluorescence staining. The brain sections from AA, SS, $SS^{FXNWT}$ and $SS^{FXNKO}$

mice were histologically assessed for markers of white matter injury using double immunofluorescence staining for non-phosphorylated antibody anti-neurofilament H (SMI32, Biolegend, cat# 801702) and myelin basic protein (MBP, Proteintech, cat# 1048-1-AP). Immunostaining was also performed to detect the axonal damage marker, the astrocyte activation marker, glial fibrillar acidic protein (GFAP; Proteintech, cat# 16825-1-AP) and the frataxin (FXN; Abcam, cat# ab197963). All images were obtained with an Olympus AX70 microscope. Staining intensity for MBP and SMI32 were determined within the corpus callosum (CC) and external capsule (EC) area by selecting a specific region using Fiji-Image J software. The mean staining intensity ratio from 6 tissue sections was used to determine the SMI32/MBP ratio for each mouse.

***Isolation of cerebral astrocytes***

Primary astrocytes were isolated from single-cell suspensions of mouse brain tissue using microbeads labeled with the astrocyte-specific anti-ACSA-1 antibody (Miltenyi Biotec; #130-095-826). The expression of frataxin in astrocytes was detected using flow cytometry.

***In vivo cognitive assessment [4]***

The $SS^{FXNWT}$ and $SS^{FXNKO}$ mice were placed in a Y-Shaped Maze to assess their spatial memory function. The number of entries and the total time spent in each arm of the maze were recorded to calculate and compare the percentage of alternations (4 trials per mouse) using the Anymaze software. The percentage of spontaneous alternations is the ratio of the number of successful alternations to the number of total alternations minus 2. Next, we performed the Novel Object Recognition (NOR) Test to evaluate short-term memory deficits. Mice were placed in the center of the arena (a plastic chamber measuring 40 cm long × 40 cm wide × 35 cm high) for 10 minutes to gain familiarity with two identical, non-appetitive objects placed 25 cm away in two corners (~8 cm from adjacent corner walls). Following the pre-test phase, mice were placed in the home cage for one hour. For the novel object test phase, one original object was replaced by a novel object (both objects were consistent in height and volume but different in shape and color), and mice were placed back into the arena for 3 minutes. Object exploration was recorded when the mice approached an object with their snout within a 2-cm perimeter around the object; the time spent exploring both objects was recorded. The exploration time (%) [(exploration time with novel object / total exploration time) × 100] and the discrimination index [(exploration time with novel object − exploration time with familiar object)/ total exploration time] were calculated [5]. One day prior to Y-maze and NOR data collection, the mice were pre-conditioned. Data were collected 3 times marked herein as Day 0, Day 7, and Day 14.

***<u>Statistical analysis</u>***

Data are presented as mean±SEM with individual data points. The data in each experimental group were considered as normally distributed based on Shapiro-Wilk test and histogram. The data were analyzed using a two-tailed, paired, or unpaired Student's t-test to determine the statistical significance between experimental groups. GraphPad Prism 10 software was used for generating the graphs and performing all statistical analyses. All p-values are reported in the figure legends. A p-value less than 0.05 was considered significant.

**References to the Supplementary materials**
